# Targeted proteomics informed by long-read RNA-sequencing reveals protein isoform expression associated with bone mineral density

**DOI:** 10.64898/2026.10.28.759629

**Authors:** Micah D. Lehe, Vasilii Pavelko, Erin D. Jeffery, Vedavathi Madhu, Larry D. Mesner, Megan D. Schertzer, Abhijit S. Dighe, Charles R. Farber, Gloria M. Sheynkman

## Abstract

The expanding catalog of sQTLs from projects like GTEx has established that genetic risk for complex traits is frequently mediated through RNA splicing. Yet it is generally unknown whether disease-associated splice events produce stable, distinct protein isoforms and whether they are functionally consequent. Even the deepest shotgun proteomics efforts only capture a minority of annotated splice events; isoform-specific peptides are systematically under-sampled by nature of being lower abundance and intrinsically harder to detect. Here, we implement a genetics-guided targeted mass spectrometry (MS) strategy to detect splice-derived protein isoforms prioritized by sQTL colocalization. Using internal standard parallel reaction monitoring (IS-PRM), we targeted 288 peptides corresponding to 166 putative BMD-associated isoforms in differentiating osteoprogenitor cells and patient-derived primary mesenchymal stem cells (MSCs). Targeted acquisition increased recovery of isoform-informative peptides relative to conventional DDA, rescued splice events that were undetected or sparsely sampled under global acquisition, and confirmed protein-level expression of genetically prioritized isoforms in clinical material. Together, this work establishes that targeted MS is a useful approach to support validation of protein expression in studies of the genetics of splicing.

## INTRODUCTION

Alternative splicing is a regulated process through which one gene produces multiple RNA and protein isoforms^1,2^. By the selective inclusion or exclusion of protein-coding regions of RNA called exons, splicing can alter the sequence of an RNA transcript and change the amino acid sequence of the expressed protein^3^. Dysregulation of splicing is pervasive in development and disease^4–6^, and growing evidence indicates that genetic risk for complex traits can be mediated through variation in RNA processing, in addition to changes to overall gene expression^7–9^.

Systematic evidence for the disease relevance of splicing has emerged from statistical genetics. Genome-wide association studies (GWAS) identify genetic variants associated with complex traits, and splicing quantitative trait locus (sQTL) analyses link genetic variants to RNA splicing events. Colocalization analyses then test whether a shared causal variant underlies both an sQTL and a disease association, prioritizing variants likely to influence disease risk through alternative splicing. Large-scale functional genomics efforts, such as the Genotype-Tissue Expression (GTEx) consortium and disease-focused sQTL studies, have linked thousands of splicing events to complex traits and diseases^7,10–12^.

One such complex trait with a significant splicing component is bone mineral density (BMD), a highly heritable trait strongly associated with osteoporosis risk^13^. In recent work, we investigated isoform regulation underlying BMD^14^. By integrating colocalized BMD sQTLs with long-read RNA-sequencing data generated from human fetal osteoblasts (hFOBs), a model of osteoblast differentiation^15^, we leveraged long-read transcriptomic data to delineate the full-length isoforms predicted to arise from BMD-associated splice events. This approach enabled isoform-level interpretation of genetic risk signals not readily resolved by short-read RNA-seq alone. Our analysis identified over 1,000 BMD-associated sQTLs in 458 protein-coding genes, nominating 2,029 candidate isoforms as putative mediators linking genetic variation to bone phenotypes.

While sQTLs and full-length RNA isoforms provide evidence of a potential role of alternative splicing in BMD, they do not confirm whether such transcripts give rise to stable protein isoforms, as changes in RNA expression do not always lead to proportionate changes in protein levels^16–18^. Direct detection of protein isoforms provides supporting evidence for whether a disease phenotype is associated with altered protein isoform abundance or, simply, reduced overall protein expression. Thus, fully characterizing alternative splicing in BMD requires detecting the predicted protein isoforms and defining their functional consequences.

Mass spectrometry (MS) remains the workhorse platform for comprehensive protein measurements. In bottom-up (shotgun) MS workflows, proteins are enzymatically digested into peptides, and evidence for isoform expression depends on detection of isoform-informative peptides that map uniquely to specific splice junctions or exons^19,20^. In practice, alternative splicing events are challenging to detect with traditional bottom-up MS. Few peptides are unique to isoforms or junctions of interest^21,22^. Approximately 20 percent of all theoretical tryptic peptides in the human proteome are isoform-specific, and the detectable fraction is likely lower in practice^22^.

Beyond peptide availability, acquisition strategy further limits detection. Data-dependent acquisition (DDA), a common mode for shotgun MS, samples precursor ions semi-stochastically based on abundance^23^. As a result, low-abundance, isoform-informative peptides—particularly those arising from sub-stoichiometric splice events—are frequently underrepresented in global acquisition workflows. Even in highly resourced efforts, such as one study that coupled multi-protease digestion and deep fractionation with DDA, amounting to over 2,400 MS injections, 26 percent of annotated splice events have been captured at the protein level^24^. This is still a minority of splice events, and the time and labor required to replicate these results is a major obstacle to widespread application.

Previously, we developed a targeted MS method capable of sample acquisition of hundreds of isoform-informative peptides in a single injection^25^. This approach adapts internal standard parallel reaction monitoring (IS-PRM)^26^ using the Tomahto acquisition framework^27^. In a prior proof-of-concept study, we utilized IS-PRM to detect protein isoforms and achieved a three-fold increase in detected isoforms compared to DDA. We therefore hypothesized that a genomics-guided targeted MS strategy—directing acquisition specifically toward splice junctions prioritized by sQTL colocalization rather than sampling the full proteome—would be well suited for validating the presence of putative protein isoforms associated with BMD.

Here, we apply a genomics-guided IS-PRM assay to detect protein isoforms associated with BMD sQTLs. First, we present a DDA-based MS analysis of mineralizing human fetal osteoblasts (hFOBs) to provide a comprehensive view of the osteoblast proteome and observe limitations of shotgun MS for isoform detection. We then report an IS-PRM assay designed to target 288 peptides from alternative protein isoforms of 36 genes with BMD-associated sQTLs. We show that IS-PRM increases detection rates and reproducibility of sQTL-prioritized, isoform-informative peptides. Finally, we apply the targeted assay to patient-derived mesenchymal stem cell populations, demonstrating feasibility of detecting genetically prioritized splice-derived isoforms in a clinical sample context.

## RESULTS

### Shotgun MS provides broad osteoblast proteome coverage but limited isoform-level resolution

To establish a proteome-wide baseline and assess isoform resolution under global acquisition, we performed shotgun MS analysis of human fetal osteoblast (hFOB) cultures undergoing mineralization over the course of ten days (**Supplemental Figure 1**). Peptides generated from each hFOB sample were labeled with isobaric tags (TMTpro), combined, separated into 21 fractions via high-performance liquid chromatography (HPLC), and analyzed via DDA on an Orbitrap Eclipse instrument^28^. To ensure that predicted splice-derived isoform sequences were included in the search space, a custom protein database predicted from hFOB full length transcripts derived from long-read RNA sequencing was used for MS searching.

In all, 7,860 proteins and 51,713 unique peptide sequences were detected (**Supplemental Figure 1**). To ask how well this captured the peptides needed for isoform-level interpretation, we compared these detected peptides to the theoretical peptide space generated by in silico tryptic digestion of the long-read–informed hFOB proteome (**Figure 1A**). Of the 678,267 theoretical peptides, only 6.8% were detected experimentally. Isoform-specific peptides (i.e., uniquely mapping to a specific transcript isoform) were particularly underrepresented: 3.1% (5,448 of 174,511) were detected. Thus, peptides that are most informative for isoform validation are disproportionately under-sampled in global DDA analyses.

**Figure 1:**
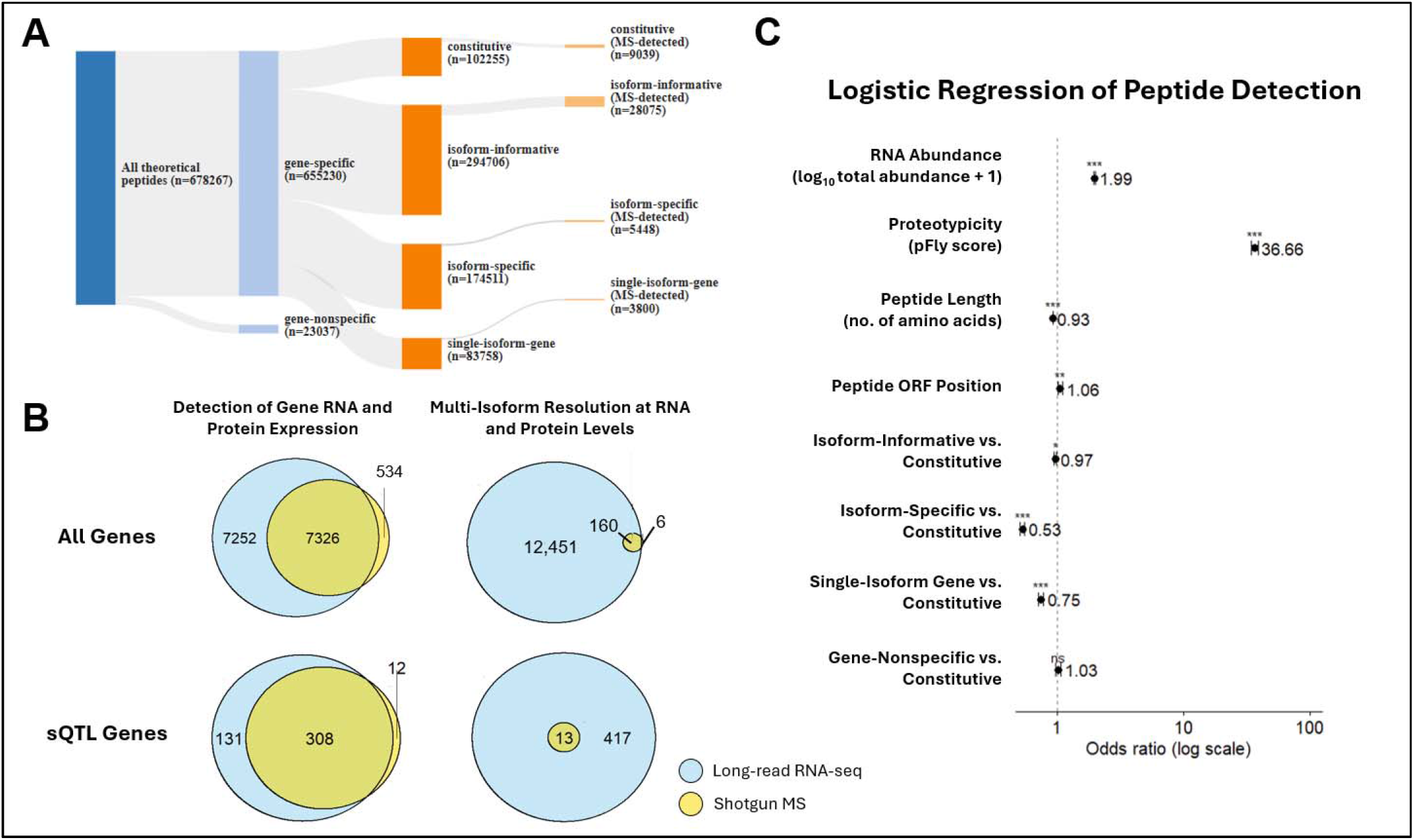
Global proteome profiling of differentiating human fetal osteoblasts (hFOBs). A) Sankey plot of all theoretical tryptic peptides generated from predicted hFOB isoforms with zero missed cleavages allowed. Peptides are classified by gene and isoform specificity and further delineated by detection with DDA-MS (21 fractions). B) Number of protein-coding genes detected (left) and number of protein-coding genes with two or more alternative isoforms detected (right) in mineralizing hFOBs with long-read RNA sequencing versus shotgun MS analysis of 21 peptide fractions. Results include all detected genes (top) and detected genes with colocalized BMD sQTLs (bottom). C) Logistic regression of detected versus undetected theoretical hFOB peptides with 21-fraction DDA-MS, with odds ratio and 95% confidence interval displayed. Significance notation is as follows: ns > 0.1, . ≤ 0.1, * ≤ 0.05, ** ≤ 0.01, *** ≤ 0.001.

The impact of limited peptide sampling extends to isoform-level inference. Although approximately half of all genes detected at the RNA level were also observed at the protein level (7,860 proteins out of 14,578 genes), evidence of more than one protein isoform per gene was rare. Only 166 genes (∼1% of detected genes) showed MS support for multiple isoforms, compared to 12,611 genes with multiple transcripts detected at the RNA level (**Figure 1B**). A similar pattern was observed among genes with colocalized sQTLs. DDA detected most sQTL-associated genes (320 proteins versus 439 genes at the RNA level) yet resolved alternative protein isoforms for only 13 of 417 genes with multiple transcripts (**Figure 1B**). Together, these results indicate that gene-level protein detection does not translate to isoform-level resolution under global acquisition.

Given the underrepresentation of isoform-informative peptides in global acquisition, we next asked whether this depletion could be explained by intrinsic peptide properties or transcript abundance, or whether isoform-informative status itself independently reduces detectability under DDA. We performed logistic regression modeling peptide detection as a function of RNA expression, isoform specificity, peptide position within the isoform ORF, peptide length, and proteotypicity (i.e. a metric of a peptide’s compatibility with MS based on physical characteristics^29^). Each theoretical tryptic peptide derived from *in silico* digestion of the long-read– informed hFOB proteome was labeled as detected or undetected in the deep-fractionation DDA dataset and included in the model.

Proteotypicity was the strongest predictor of detection, increasing a peptide’s odds of detection by about 37-fold per one-unit score increase (**Figure 1C**). RNA presence and abundance were also positively associated with detection. Notably, isoform-specific peptides were approximately half as likely to be detectable as constitutive peptides shared across all isoforms of the same gene, even after accounting for these other features. Together, these findings indicate that intrinsic peptide properties and transcript abundance influence detection probability under DDA, and that isoform-informative peptides remain less likely to be detected even after adjusting for these variables. However, this association does not exclude the possibility that some splice-derived transcripts may not yield stable or translated protein products, underscoring the importance of sensitive and isoform-resolved protein-level validation.

### IS-PRM increases detection of BMD-associated isoform-informative peptides in differentiating osteoblasts

To address the issue of low isoform coverage observed under global acquisition, we employed IS-PRM, which focuses MS acquisition on a pre-specified set of peptides. We used this approach to target BMD-associated isoforms previously nominated through sQTL colocalization and long-read transcriptome analysis (**Figure 2A**). We first developed a prioritization strategy, building the list hierarchically from genes and down to individual peptides. At the gene level, candidate peptides were drawn from loci harboring BMD-colocalized sQTLs. At the isoform level, target transcripts were prioritized based on the strength of sQTL colocalization with BMD (H4 posterior probability) and RNA abundance (counts per million, CPM) (**Supplemental Figure 2, Methods**). At the peptide level, we selected isoform-informative peptides uniquely distinguishing predicted alternative splice variants. Constitutive peptides shared across all isoforms of the same gene serve as gene-level reference measurements. Final peptide selection additionally considered standard LC–MS constraints, including peptide length (7–21 amino acids) and predicted retention time (RT) within the chromatographic gradient. In total, we selected 288 target peptides corresponding to 166 predicted protein isoforms across 36 protein-coding genes that had BMD colocalizations.

**Figure 2:**
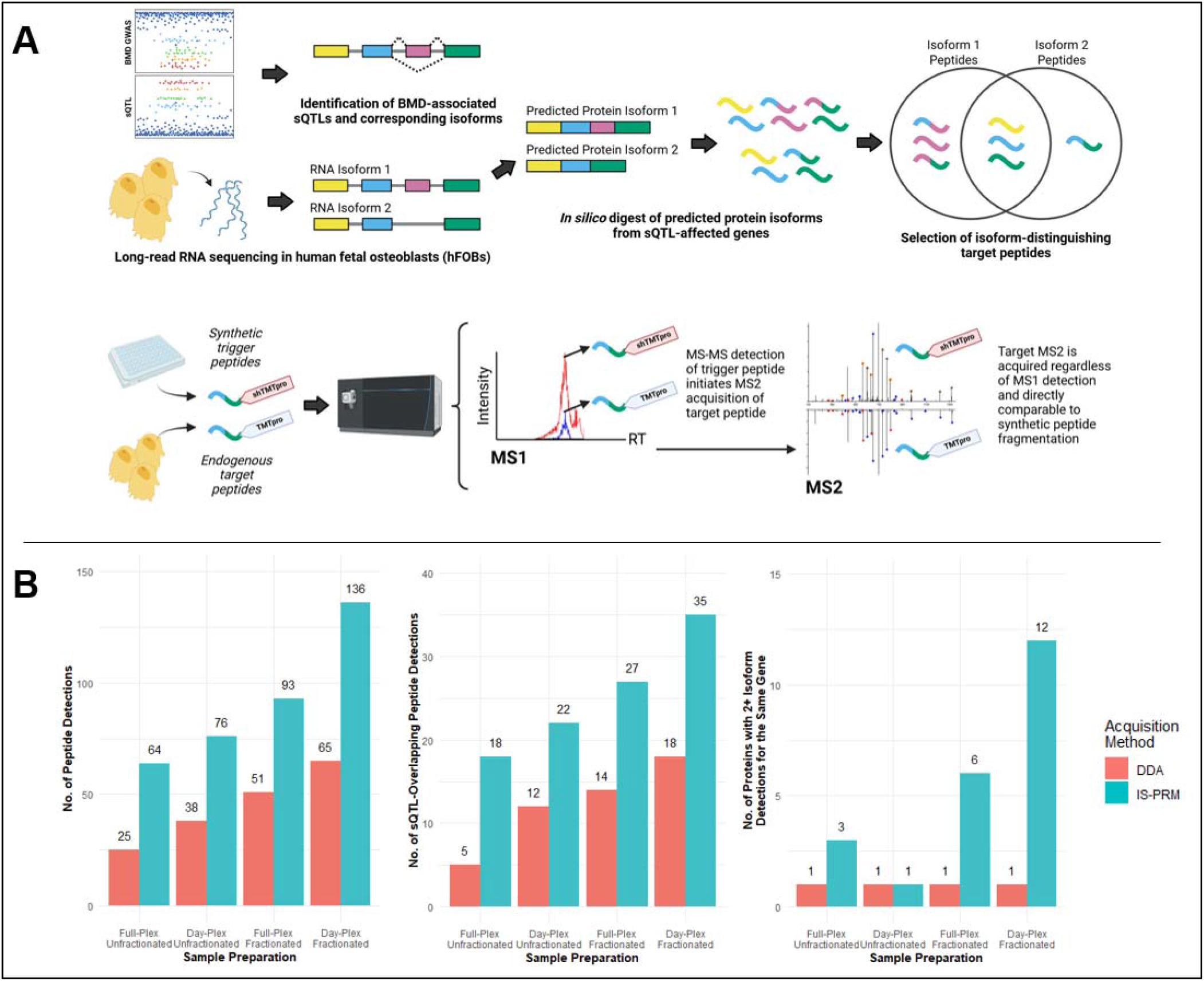
Increased detection of splice-specific sQTL peptides with IS-PRM. A) Schematic of selecting isoform-specific or splice event-specific peptides corresponding to BMD sQTLs and transcripts expressed in the osteoblast model based on long-read RNA sequencing data. Target peptides are detected via IS-PRM as described in Yu et al.^27^ and Korchak et al.^25^. B) Bar plot of numbers of detected target peptides, detected targets spanning an sQTL junction, and genes with multiple resolved protein isoforms, comparing IS-PRM versus a standard DDA approach. Sample fractionation and TMTpro multiplexing are indicated on the x-axis (see Supplemental Figure 3). A multiple isoform detection event was determined for a given protein when two or more mutually exclusive peptides (i.e., two or more peptides generated by different alternative splicing events) were detected within the same peptide sample.

To test whether IS-PRM would recover these target peptides more effectively than global acquisition, we benchmarked the two approaches on matched hFOB samples. IS-PRM relies on synthetic "trigger" peptides: heavy-labeled versions of each target peptide are spiked into the sample, and their detection in MS1 triggers targeted fragmentation of the corresponding co-eluting endogenous peptide. We synthesized trigger peptides for all 288 targeted sequences and labeled them with super heavy TMTpro (shTMTpro), a TMT variant that introduces a precursor mass offset to distinguish trigger peptides from endogenous peptides (**Supplemental Figure 3, Methods**). For the biological samples, we prepared hFOB cultures across 0–6 days of differentiation (day 0, 2, 4, and 6; n=3 per timepoint; 12 samples total), labeled each with a unique TMTpro reagent, and spiked the pooled trigger mixture into the multiplexed sample prior to analysis.

To evaluate IS-PRM across the range of conditions a real experiment might encounter, we varied two parameters that together define how much instrument time and coverage an MS experiment demands: sample complexity (via multiplexing) and fractionation depth (**Supplemental Figure 3**). For multiplexing, we compared a full-plex sample in which all 12 TMTpro-labeled samples were pooled into a single high-complexity mixture— against day-separated plexes, in which only the three biological replicates from each timepoint were pooled.

For fractionation, each multiplexed sample was analyzed either directly (unfractionated) or after high-pH reverse-phase fractionation into multiple peptide fractions. Every preparation condition was acquired using both DDA and IS-PRM, allowing comparison of target peptide coverage and isoform resolution between the two acquisition modes.

IS-PRM consistently outperformed DDA across every preparation condition tested. On average, IS-PRM achieved approximately two-fold greater coverage of target peptides relative to DDA, and this increase held for the subset of peptides overlapping BMD sQTL junctions (**Figure 2B**). Under the conditions maximizing acquisition depth (day-separated three-plex with fractionation), IS-PRM detected 138 of the 288 target peptides, compared to 65 with DDA. Additionally, IS-PRM detected 35 sQTL junction-spanning peptides (out of 104 total) under the same conditions. The difference was even greater at the isoform level. DDA resolved multiple isoforms for only one target protein, whereas IS-PRM resolved multiple isoforms for 12 genes with BMD-colocalized sQTLs. Directing acquisition towards genetically prioritized splice events therefore substantially improves isoform-level resolution relative to global proteome sampling.

### IS-PRM improves detection and reproducibility of BMD-associated splice-specific peptides

To illustrate how IS-PRM improves both the detection and reproducibility of BMD-associated splice events relative to DDA, we examined specific cases in detail. In several instances, IS-PRM detected splice events that DDA missed entirely. *TRIP6* (thyroid receptor-interacting protein 6) is involved in cytoskeletal reorganization and focal adhesion regulation^30^ and its expression is elevated in osteosarcoma^31^. We previously identified a BMD-associated sQTL at the *TRIP6* locus and, by combining RNA sequencing with open reading frame prediction, predicted that the event produces a frameshift near the start codon, giving rise to an alternative isoform with a C-terminal sequence entirely distinct from the canonical isoform^14^ (**Figure 3A**). However, no DDA method, including deepest 21 fraction analysis, detected peptides specific to either predicted splice junction. IS-PRM, in contrast, detected a junction-specific peptide from each isoform, despite a 50-fold difference in their RNA abundances (6 CPM vs. 348 CPM).

**Figure 3:**
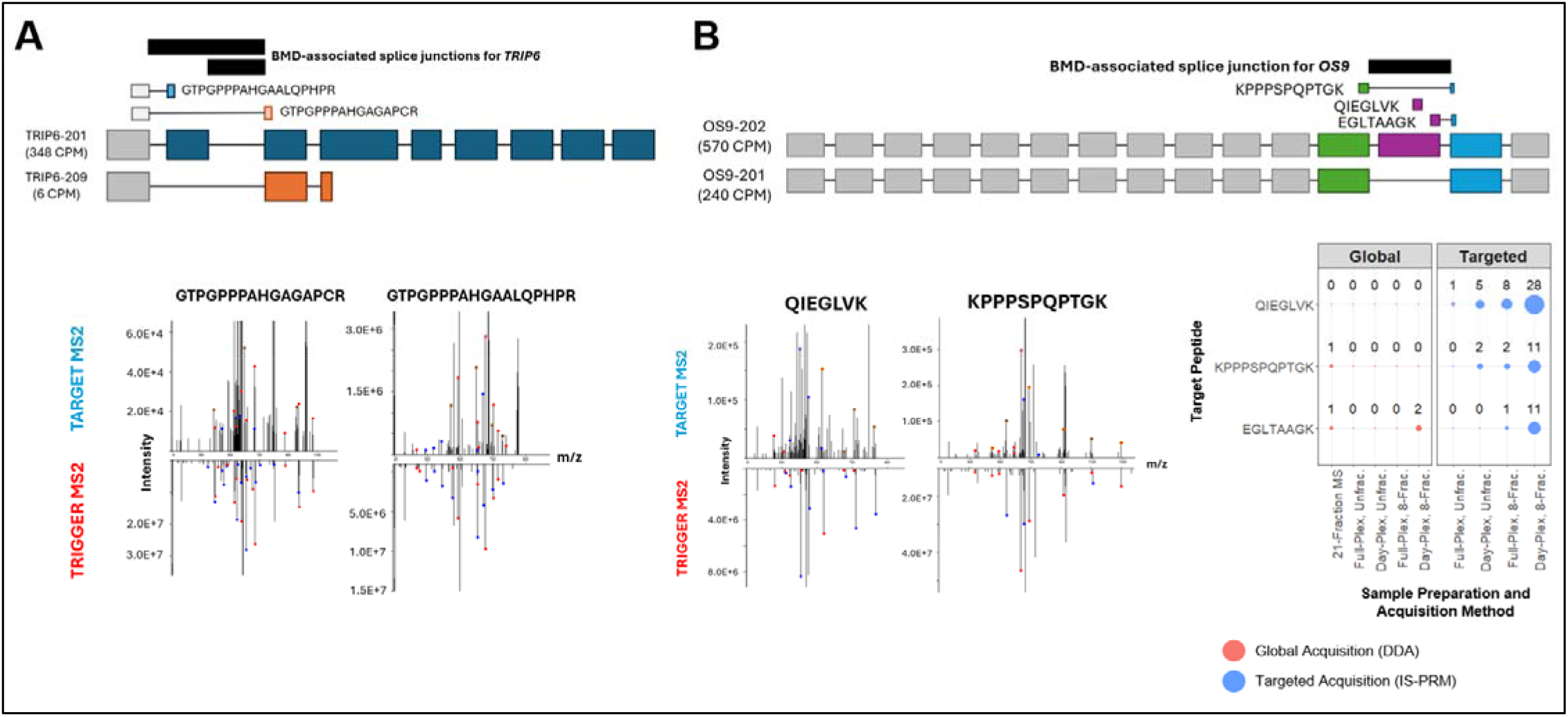
Resolution of BMD-associated splice event in TRIP6 and OS9 protein isoforms. A) Schematic of two TRIP6 isoforms and a pair of mutually exclusive target peptides overlapping with a BMD-associated splice junction. Bottom: PSM mirror plots depicting target peptide MS2 spectra and the immediately preceding trigger peptide MS2 spectra. B) Schematic of two major OS9 isoforms predicted from long-read RNA sequencing with a BMD sQTL splice junction (black) corresponding to exclusion of an exon (purple) near the C-terminus. Target peptides detected with IS-PRM are aligned with the protein isoform schematics. Bottom left: PSM mirror plots depicting target peptide MS2 spectra and the immediately preceding trigger peptide MS2 spectra. Bottom right: dot plot of total peptide spectral matches (PSMs) for OS9 target peptides using DDA (red) and IS-PRM (blue). Fractionation and multiplexing strategies employed for DDA and IS-PRM acquisition are noted on the x-axis (see Supplemental Figure 3).

Even when DDA did detect splice-specific peptides, detection was often too sparse to provide reliable evidence of isoform expression across conditions. As an example, *OS9* (endoplasmic reticulum lectin) is a regulator of ER-associated degradation^32,33^ and is highly upregulated in osteosarcomas^34^. The BMD-associated sQTL at *OS9* involves skipping of a 54-amino acid exon near the C-terminus (**Figure 3B**). DDA did detect peptides diagnostic of both the exon-skipping (KPPPSPQPTGK) and exon-inclusion (EGLTAAGK) events, but only marginally: the deepest DDA condition (21 fractions) yielded just one peptide spectral match (PSM) per peptide across all fractions, and in the matched side-by-side comparison with IS-PRM, DDA produced only two PSMs for EGLTAAGK and none for any other splice-specific peptides. IS-PRM, under the same matched conditions, yielded up to 11 PSMs for KPPPSPQPTGK and EGLTAAGK and additionally detected 28 PSMs for a third exon-inclusion peptide (QIEGLVK) that was absent from all DDA analyses. In this case study, splice events sitting at the margin of detectability under DDA, regardless of fractionation depth, therefore became reproducibly and confidently measured with IS-PRM.

### IS-PRM reveals quantitative RNA-protein correlation for BMD-associated TPM2 isoforms

The preceding examples demonstrate that IS-PRM improves qualitative detection of splice-specific peptides. We next asked whether IS-PRM could capture quantitative isoform dynamics across osteoblast differentiation. We focused on tropomyosin-2 (TPM2)^35^, for which we previously identified a BMD-associated exon switching event with direct functional role in mineralization^14^. Selective knockdowns of Exon 6- or Exon 7-containing transcripts produce opposing effects on bone mineralization phenotype^14^. At the RNA level, pre-mineralization osteoblasts preferentially express *TPM2* transcripts containing Exon 7, but the dominant transcripts shift to those containing Exon 6 at the onset of mineralization (**Figure 4B**). Our IS-PRM analysis recapitulated this switch at the protein-level, with target peptides specific to Exon 7 decreasing in abundance over the course of hFOB differentiation (**Figure 4C**). Peptides specific to Exon 6 as well as constitutive peptides shared by all TPM2 isoforms showed a slight upward trend. IS-PRM thus captured the expected directional shift in TPM2 isoform usage at the protein level, with the clearest signal corresponding to the decline of Exon 7-containing isoforms. This result demonstrates a proof-of-concept that targeted acquisition can move beyond binary isoform detection to reveal quantitative isoform dynamics at the protein level.

**Figure 4:**
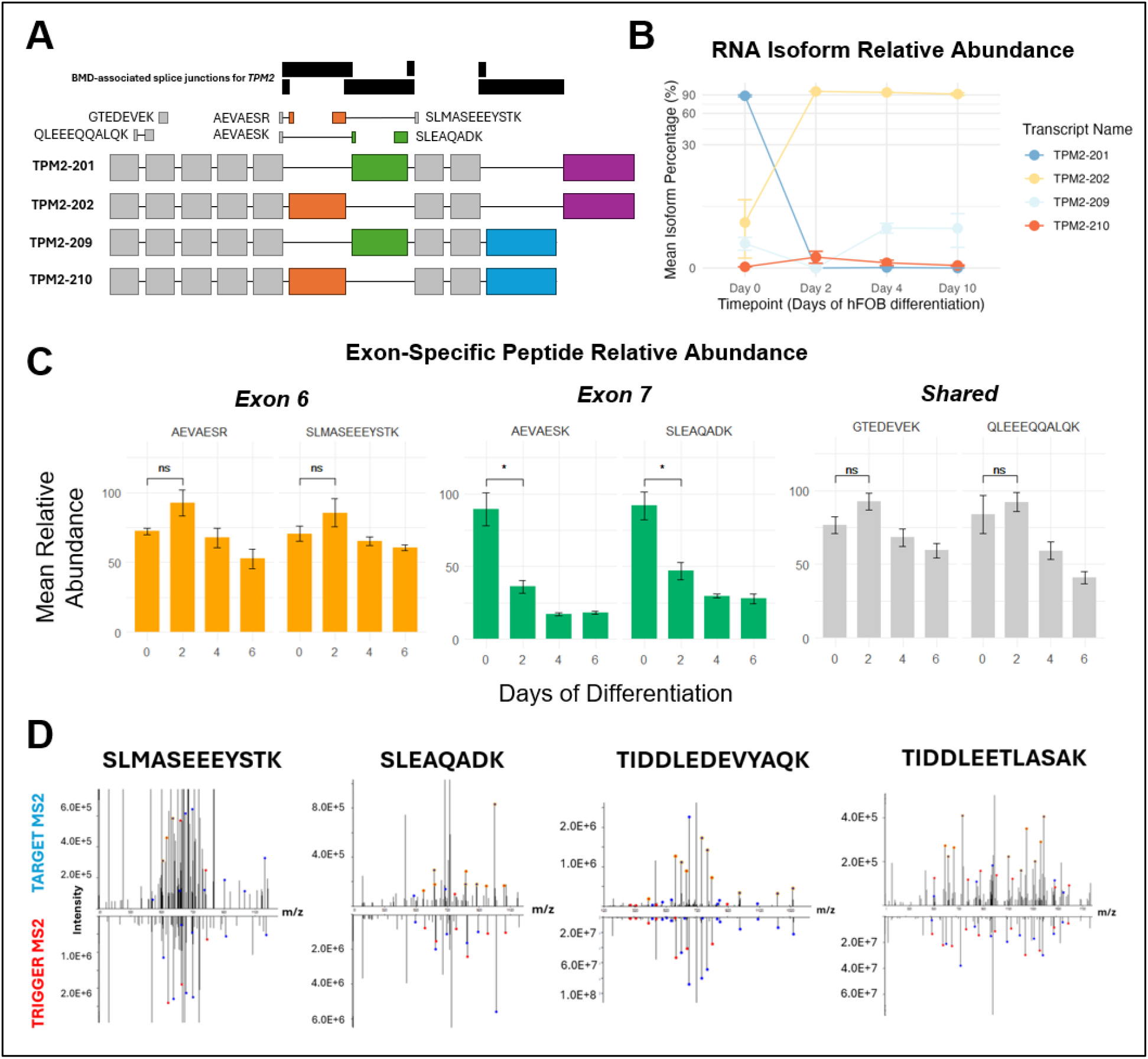
RNA-protein correlation of TPM2 isoform expression in mineralizing hFOBs. A) Schematic of the four major TPM2 protein isoforms generated by BMD sQTLs (black). The isoforms contain one of two exons in the central portion of the protein (orange and green) and one of two C-terminal domains (blue and purple). Target peptides detected with IS-PRM are aligned above the isoforms. B) Differential usage of each *TPM2* mRNA across hFOB differentiation. Each isoform’s fractional contribution to total *TPM2* expression was determined from long-read RNA sequencing. C) Mean relative abundance of TPM2 target peptides measured with IS-PRM over six days of hFOB mineralization. TMTpro reporter ion abundances for each peptide spectral match (PSM) collected with IS-PRM were scaled 0 to 100. Error bars represent standard deviation of the mean relative abundance for each biological replicate for each day of mineralization. Hypothesis testing was conducted using a standard t-test. Notation for p-values is as follows: ns > 0.05, * ≤ 0.05, ** ≤ 0.01, *** ≤ 0.001, **** ≤ 0.0001. Bars denote standard deviation. D) PSM mirror plots depicting target peptide MS2 spectra (top) and the immediately preceding trigger peptide MS2 spectra (bottom).

### RNA abundance is the strongest predictor of target peptide detectability

Even under the deepest IS-PRM condition, 150 of the 288 target peptides remained undetected. Given that this represents about half of the targets, we asked what peptide and transcript features distinguished detected from undetected targets. Similar to Figure 1C, we used logistic regression to model peptide detection status as a function of RNA abundance, proteotypicity, and isoform specificity (**Figure 5A**). Proteotypicity scores were modestly but significantly higher for detected versus undetected peptides (p ≤ 0.01), although the distributions largely overlapped (**Figure 5B**), indicating that intrinsic peptide detectability still influences detectability even under targeted acquisition. RNA-based metrics, however, provided stronger discrimination. For each target peptide, we calculated the total expression of all transcripts encoding that peptide and the isoform ratio, defined as the fraction of the gene’s total expression contributed by transcripts containing the peptide’s coding region (**Figure 5C**). Both total RNA expression (**Figure 5D**) and isoform ratio (**Figure 5E**) were significantly higher for detected peptides (p ≤ 0.0001). These results indicate that while proteotypicity imposes a secondary constraint on detection even with targeted acquisition, RNA expression—both absolute and isoform-fractional—is the more reliable predictor of whether a splice-specific peptide will be captured by IS-PRM. These results suggest that once sampling bias is eliminated through targeted acquisition, protein-level isoform detection is strongly dependent on the underlying transcript expression.

**Figure 5:**
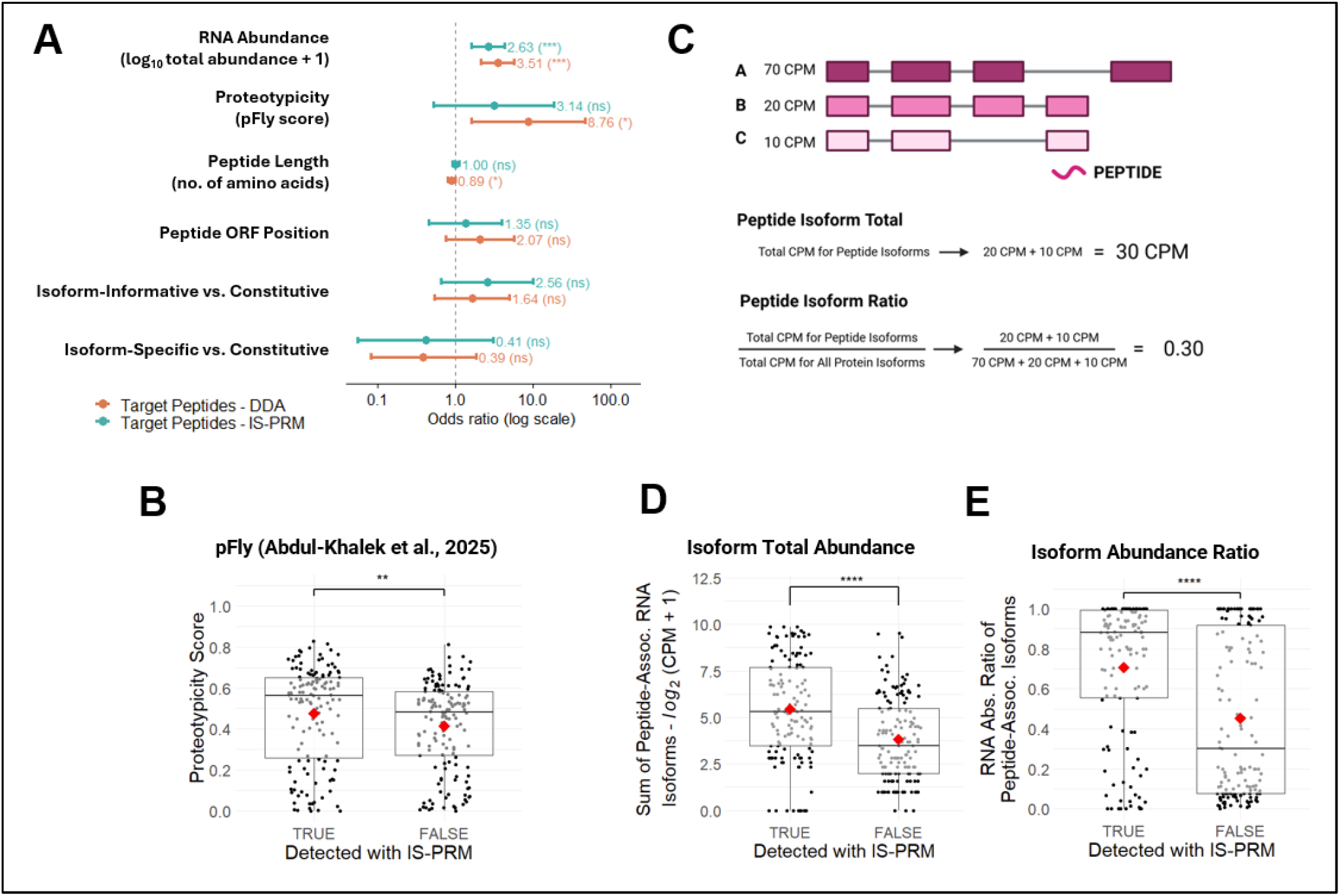
Analysis of detectability predictors for BMD target peptides with IS-PRM. A) Logistic regression of detected versus undetected BMD target peptides in IS-PRM and DDA (conducted in parallel with IS-PRM, not 21-fraction MS), with odds ratio and 95% confidence interval displayed. Significance notation is as follows: ns > 0.1, . ≤ 0.1, * ≤ 0.05, ** ≤ 0.01, *** ≤ 0.001. B) Comparison of proteotypicity scores for target peptides using the pFly model^44^ based on amino acid sequence and protease efficiency. Hypothesis testing was conducted using Wilcoxon rank-sum test. C) Schematic of calculating a total RNA abundance and RNA isoform ratio for each target peptide. D) Comparison of target peptides by total RNA isoform abundance. Hypothesis testing was conducted using Wilcoxon rank-sum test. E) Comparison of target peptides by RNA isoform ratio. Hypothesis testing was conducted using Wilcoxon rank-sum test. For box plot statistical tests, p-value notation is as follows: ns > 0.05, * ≤ 0.05, ** ≤ 0.01, *** ≤ 0.001, **** ≤ 0.0001. Mean values in each boxplot are represented by a red diamond.

### IS-PRM application to human MSCs validates BMD-relevant alternative splicing events in the proteome

Having established IS-PRM performance in the hFOB model, we next asked whether the same BMD-relevant isoforms could be detected using our assay in the analysis of patient-derived samples and thus providing support for isoform protein expression in a clinically relevant context. We isolated cells from the bone tissue of four patients undergoing hip replacement and cultured them in three conditions of increasing degree of mineralization (**Figure 6A**). The resulting 12 samples (4 patients, 3 conditions) were TMT-labeled and multiplexed as in the hFOB experiments. The assay detected 129 target peptides in patient samples, 109 of which were also observed in hFOBs (**Figure 6B**). Among 31 detected peptides that overlapped with a BMD splice junction (i.e., an sQTL), 28 were also detected in hFOBs. Eight genes exhibited evidence of two or more stable protein isoforms in patient samples, including seven that were also resolved in the hFOB model. Based on these results, detection performance of the BMD IS-PRM assay transfers reproducibly from the cell-line model to patient-derived clinical material.

**Figure 6:**
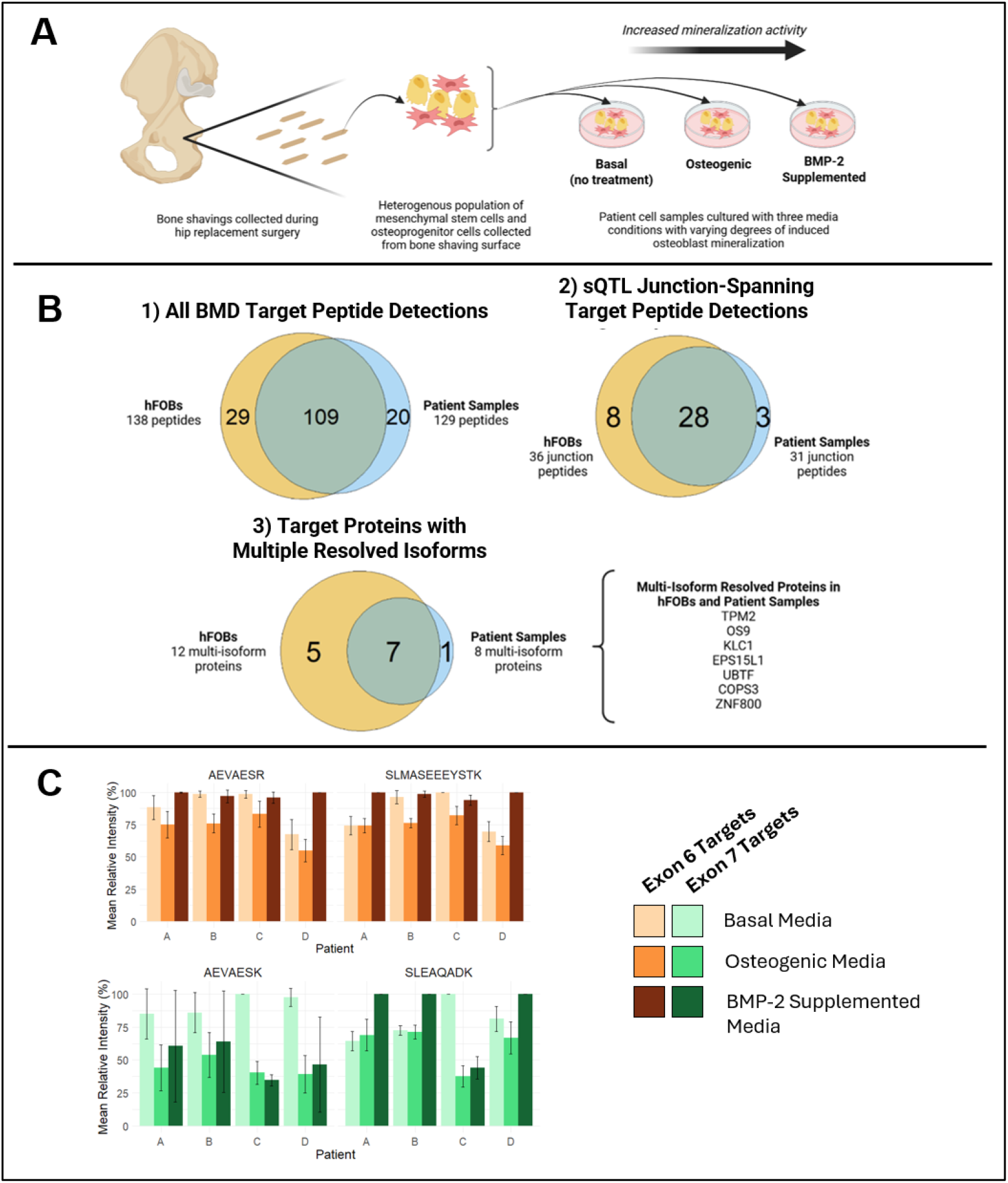
Detection of BMD-associated isoforms in patient-derived bone cell populations. A) Schematic of isolation of bone cells from patient bone shavings. Cell cultures consisting of mesenchymal stem cells (MSCs), committed osteoprogenitors, and mature osteoblasts and osteoclasts were grown in three media conditions (with varying effects on osteoblast commitment and differentiation) for 10 days prior to MS analysis. B) Venn diagrams comparing BMD target peptide and isoform detections between hFOB model and patient samples. Proteins with successful multi-isoform resolution in both hFOBs and patient samples are listed. C) Relative abundance of select TPM2 target peptides in clinical samples, distinguished by patient donor and media condition.

We next asked whether the quantitative TPM2 exon-switching dynamics observed in hFOBs (**Figure 4C**) were recapitulated in the patient-derived samples (**Figure 6C**). One target peptide (AEVAESK) somewhat followed the expected trend, but most TPM2 target peptides showed no consistent directional pattern across the patient samples. Several factors may contribute, though we cannot distinguish among them with the current data.

First, patient-derived cultures are compositionally heterogeneous, mixing osteoblasts with other bone-resident cell types that may express TPM2 isoforms at different levels. Second, TPM2 isoform diversity in primary tissue may extend beyond the canonical exon 6/7 switch that dominates the hFOB model. Nevertheless, these observations indicate that while detection-level reproducibility transfers cleanly to clinical samples, quantitative isoform dynamics observed in a homogeneous cell line may not generalize directly to patient material without more defined sample inputs, such as sorted osteoblast populations.

## DISCUSSION

Genome-wide association studies and sQTL analyses have nominated thousands of splice events as potential mediators of genetic risk for complex traits, yet systematic protein-level confirmation of these events has been largely absent. In this study, we demonstrate a genomics-guided targeted mass spectrometry strategy that directs acquisition specifically toward splice junction-overlapping peptides prioritized by sQTL colocalization. Using IS-PRM, we approximately doubled detection of BMD-associated isoform-informative peptides relative to conventional DDA, recovered splice events that were missed or only marginally sampled by global acquisition (e.g., TRIP6 and OS9), and demonstrated feasibility of detecting prioritized isoforms in patient-derived bone samples. Together, these findings lay the groundwork for a scalable framework for translating transcript-level sQTL predictions into protein-level confirmation.

A core feature of genomics-guided targeted proteomics is how targets are chosen. In contrast to conventional targeted proteomics, where peptides are drawn from well-characterized proteins, our starting point is a set of splice junctions nominated by statistical genetics, which must then be translated, through multiple layers of inference, into analytically tractable protein targets. Previously, we addressed the first part of this chain by integrating sQTL data with long-read RNA sequencing, defining putative protein isoforms connected to individual splicing events^14,36^. With such isoform models in hand, target selection required navigating the two intersecting constraints of biological informativeness and analytical tractability. We prioritized peptides not merely for isoform specificity within a gene, but for their ability to distinguish sQTL-associated splice events, incorporating both junction-specific peptides and shared peptides near each sQTL to provide gene-level reference measurements. Candidate peptides were further filtered according to standard MS constraints, including peptide length, predicted retention time, and proteotypicity. This multi-criteria selection process— spanning genetic prioritization, isoform-resolved protein prediction, and analytical feasibility—represents a distinct category of assay design for targeted proteomics.

Even with careful target design, genomics-guided targeted proteomics can ultimately only increase detection within the subset of peptides that are theoretically amenable to MS analysis. Some splice events do not generate analytically tractable tryptic peptides, and not all sQTL-associated genes in our study yielded targetable peptides. A second limit applies to events that do in fact yield tractable peptides, which is that detection is further dictated by intrinsic sequence properties and transcript abundance, as our regression analysis confirmed. An implication worth considering is that non-detection of peptides may carry biological meaning rather than being a mere technical limitation. For a target with adequate RNA expression, favorable peptide chemistry, and no PTM-related confounders, failure to detect the corresponding protein could reflect post-transcriptional regulation, protein instability, or nonsense-mediated decay of the underlying transcript, mechanisms that explain lack of correlation between RNA and protein levels. As genomics-guided targeted proteomics matures, negative results could become a distinct source of biological information.

Beyond binary detection, the quantitative tracking of isoform dynamics across conditions remains a challenge. TPM2 provided one example in which splice-dependent peptide abundance tracked RNA-level isoform switching. However, we found that other isoforms did not show as clear a correlation between RNA-level isoform expression and target peptide abundance (**Supplemental Figure 4**). Two explanations are plausible. First, peptides that are harder to detect will yield noisier quantitative measurements, so technical factors like proteotypicity and PTM susceptibility carry over from detection into quantification. Alternatively, the discordance may reflect real biology, as RNA abundance does not scale directly to protein expression^16–18^, and non-canonical isoform transcripts are especially prone to nonsense-mediated decay^37^, both of which would result in non-concordance between RNA and protein levels.

Several directions will extend this work. Methodologically, the synthetic trigger peptides used here provided a gold-standard spectral reference for confirming isoform-informative peptides. Such high-confidence validation will become increasingly important as expanding human genetic studies reveal rare, individualized, and disease-subtype–restricted splice variants that lack representation in public spectral libraries. However, synthetic peptide panels are costly to produce at scale. Emerging trigger-free targeting methods can achieve a similar throughput to IS-PRM by utilizing experimentally derived RT and spectral libraries^38–40^, newer iterations can target novel peptides using predicted libraries^41^, and hybrid PRM/data-independent acquisition (DIA) modes combine global and targeted analysis within a single run^42^. Clinically, the application of our IS-PRM assay to patient-derived bone samples establishes proof-of-concept that genetically prioritized protein isoforms can be detected in clinical material. Extending this toward mechanistic conclusions about the role of specific isoforms in BMD will require larger cohorts with cellular resolution to address the heterogeneity of primary bone tissue, quantitative assays capable of measuring isoform stoichiometry, and individuals with known genotypes at the relevant sQTL loci. More broadly, the strategy demonstrated here provides the protein-level validation component of a generalizable framework for connecting genetic variation to proteoform biology across complex traits and diseases.

## ACKNOWLEDGEMENTS

Research by M.D.L. was supported by the National Institute of General Medical Sciences of the National Institutes of Health under Award Number 2T32GM136615-06. Additional NIH support was provided to G.M.S. by NIGMS under Award Number 5R35GM142647-04 and the National Library of Medicine under Award Number 5R01LM014017-04. Support for A.S.D. and C.R.F. was provided by the National Institute of Arthritis and Musculoskeletal and Skin Diseases of the NIH under Award Number 1R01AR086161-01. The content is solely the responsibility of the authors and does not necessarily represent the official views of the NIH.

## AUTHOR CONTRIBUTIONS

M.D.L.: experimental design, sample preparation, data collection and analysis, manuscript writing and review. V.P.: sample preparation (hFOB) and data collection support. E.D.J.: proteomics data collection support and manuscript review. V.M.: sample preparation (clinical) and data analysis support. L.D.M.: sample preparation support (hFOB). M.D.S.: data analysis support and manuscript review. A.D.S.: sample preparation support (clinical). C.R.F.: genomics project guidance and primary supervisor for V.M., L.D.M., and A.D.S. G.M.S.: overall project conception and guidance, primary supervisor for M.D.L., V.P., E.D.J., and M.D.S.

## SUPPORTING INFORMATION

Supplemental text and figures are provided as an attached document. Supplemental tables are provided as an attached Excel document.

Table S1: Guide to TMTpro reporter ions and corresponding samples in hFOB and clinical proteomics analyses

Table S2: BMD target isoforms by PacBio (PB) accession

Table S3: BMD target peptides with isoform assignments and sQTL information

Table S4: BMD target peptide detection summary in hFOB samples

Table S5: BMD target peptide detection summary in patient-derived samples

Table S6: BMD trigger peptide detection in false-positive control experiment

Table S7: Guide to MS raw file names

## METHODS

### BMD Genomics and Long-Read Proteogenomics

Analysis of genomic and transcriptomic data used for target peptide selection is described in Abood et al^14^. Briefly, genome-wide association study (GWAS) summary statistics from the Genetic Factors for Osteoporosis Consortium (GEFOS) were used to identify bone mineral density (BMD)-associated single nucleotide polymorphisms (SNPs). Summary statistics for splice quantitative trait loci (sQTLs) across multiple tissue types were collected from the Genotype-Tissue Expression (GTEx) Portal (v.8). Potential BMD-associated splice junctions were identified by colocalizing genome coordinates for BMD GWAS SNPs with GTEx sQTLs. For every splice junction within 200 kilobases of a gene with a BMD GWAS SNP, H4 posterior probability (H4PP) was calculated to determine the likelihood that the SNP and sQTL (or splice event) shared a causal variant. sQTLs with an H4PP of ≥0.75 were considered to be the most significant predicted BMD-associated splice events.

The long-read proteogenomics (LRP) bioinformatics pipeline described in Miller et al.^36^ was used to generate a hybrid bone proteomic database. Briefly, long-read RNA sequencing reads from hFOBs (collected at 0, 2, 4, and 10 days of differentiation) were converted into proteoforms via open reading frame (ORF) prediction.

Proteoforms that matched an existing protein sequence in the GENCODE human proteome database were matched with their standardized GENCODE protein and isoform accession, while novel proteoforms were designated with their PacBio RNA-seq identification number. A hybrid FASTA file was constructed by compiling predicted hFOB proteoforms with all GENCODE human protein sequences that were not represented in the RNA-seq data. Significant BMD sQTLs identified in genomic analysis were visualized alongside long-read RNA-seq reads on the UCSC Genome Browser^45^. Isoforms were color-coded by relative transcript abundance.

Theoretical hFOB peptides were generated using the cleaver R package from Bioconductor^46,47^. Digestion parameters were set to trypsin (C-terminal digestion at lysine and arginine residues), up to one missed cleavage allowed, and peptide length between 6 and 51 amino acids long.

### BMD Target Peptide Selection

Gene candidates with significant predicted BMD-associated splice events were narrowed by considering the RNA abundance of alternative isoforms based on counts per million (CPM), the location of the sQTL(s) in the protein-coding region, and the ability to distinguish theoretical proteins isoforms with MS-compatible tryptic peptides. Once selected, protein isoform candidates were digested *in silico* into tryptic peptides using the cleaver R package from Bioconductor^46,47^. Peptides were then mapped back to the genome browser and aligned with RNA isoform reads using PoGo^48^. Candidate target peptides were selected based on their ability to distinguish between BMD-associated splice events or serve as constitutive (shared) peptides mapping to all protein isoforms in consideration for a given gene. Final target peptides were selected based on their uniqueness to their respective protein-coding genes, their size (between 6 and 21 amino acids), and their predicted retention time on the MS liquid chromatography gradient as calculated by AutoRT, a retention time prediction model^49^.

### Trigger Peptide Reconstitution

Synthetic peptides (SpikeTides™) were acquired from JPT in three sets of 96, for a total of 288 trigger peptides with matching sequences to selected target peptides. Peptides were reconstituted in 80% 0.1M ammonium bicarbonate, 20% acetonitrile. For each of the three plates, all 96 trigger peptides were pooled together at equal volume. Following concentration measurement with NanoDrop, peptides were dried via speed vac and reconstituted in 0.1% formic acid to 1 µg/µL. A 1 µg peptide aliquot then analyzed using a 120-minute DDA-MS method. Detection and approximate intensity of each trigger peptide was estimated via analysis in Skyline (version 24.1.0.414). Based on this analysis, synthetic peptides were re-pooled at varying concentration factors to improve precursor intensity of low-abundance or undetected peptides.

### Trigger Peptide shTMTpro Labeling

Following MS analysis of precursor intensities, the optimized pool of synthetic trigger peptides was labeled with super-heavy TMTpro reagent (Thermo Scientific A52040). 100 µg of peptides from the optimized pool were dried via speed vac and reconstituted in 100 µL 0.2 MM EPPS buffer (pH 8). shTMTpro (0.5 mg) reagent was reconstituted in 20 µL anhydrous acetonitrile and transferred to the 100 µL synthetic peptide solution. Labeling was conducted at room temperature for one hour with brief vortexing every 10 minutes. A 4 µL aliquot was taken for a labeling efficiency check, while the remaining reaction mixture was temporarily stored at -80°C. The label check aliquot was desalted, dried via speed vac, resuspended in 0.1% formic acid, and injected on a 120-minute DDA-MS method. The results from the MS run were analyzed with both Skyline and Proteome Discoverer to ensure a labeling efficiency rate of at least 90%. Following analysis, the reaction mixture was thawed and re-labeled with a second shTMTpro aliquot if necessary. Labeling was quenched by adding 5% hydroxylamine to the reaction mixture, vortexing, and incubating at room temperature for 15 minutes. Labeled peptides were then dried via speed vac and reconstituted in 100 µL 0.1% formic acid. Half (50 µL) of each labeled peptide pool was desalted, while the remaining half was kept for fractionation.

### hFOB Cell Cultures

Protocols for human fetal osteoblast (hFOB) (American Type Culture Center [ATCC], Manassas, VA; CRL-11372) cultures were adapted from previously described methods^14^. Briefly, twelve hFOB cultures were prepared with growth media (MEM with 10% fetal bovine serum, 1% Glutamax, 1% Pen-Strep) and incubated at 34°C, with media replaced every three days as needed. Pellets were collected from three of the cultures once all culture plates reached 80% confluence, representing hFOBs at zero days of mineralization. Growth media in all other cultures was replaced with differentiation media (MEM alpha with 10% FBS, 1% Glutamax, 1% Pen-Strep, 50 µg/µL ascorbic acid, 10 mM beta-glycerophosphate, 10 nM dexamethasone) and incubated at 39.5°C. Pellets were harvested from three of the remaining hFOB cultures following two, four, and six days of mineralization. Osteoblast mineralization was tracked alongside cell pellet collection in separate hFOB cultures via an adapted Alizarin Red staining protocol.

### hFOB Cell Lysis

Twelve hFOB cell pellets, representing four mineralization time points with three biological replicates each, were each resuspended in cell lysis buffer (6% SDS, 150 mM DTT, 75 mM Tris-HCl) and lysed with a probe sonicator set to 20 amplitude for 3-5 minutes. Lysates were heated at 95°C for 3 minutes, and protein concentration was measured via a 660nm colorimetric assay (Thermo Scientific 22660 and 22663).

### hFOB Protein Digestion

Protein lysates (100 µg of each time point and bio-replicate) were digested into peptides via filter-assisted sample preparation (FASP) according to manufacturer protocol (Millipore UFC503096). Proteins were digested on filter columns for 18 hours at 37°C in 50 mM ammonium bicarbonate with 1 µg trypsin per 100 µg hFOB protein. Following digestion, peptides were eluted via centrifugation and concentration was measured via NanoDrop (Protein A280).

### hFOB Peptide TMTpro Labeling

The full elution volume from each hFOB peptide FASP was dried via speed vac and reconstituted in 100 µL 0.2 MM EPPS buffer (pH 8). Each of the twelve hFOB peptide samples were assigned a unique TMTpro reagent (**Supplementary Table 1**). A separate peptide sample collected from a Jurkat cell culture was included and assigned a unique TMTpro 16-plex reagent (Thermo Scientific A44522). A 0.5 mg aliquot of each selected TMTpro label was resuspended in 20 µL anhydrous acetonitrile and transferred to its respective peptide sample. Labeling reactions were conducted at room temperature for one hour with brief vortexing every 10 minutes. After labeling, a 2 µL aliquot of each reaction was taken for a labeling efficiency check; the remaining reaction mixtures were temporarily stored at -80°C. The label check aliquots were pooled together into one multiplexed sample, desalted, dried via speed vac, resuspended in 10 µL 0.1% formic acid, and injected on a 120-minute DDA-MS method. The results from the MS run were analyzed with Proteome Discoverer to ensure a labeling efficiency rate of at least 90%. Labeling was quenched by adding 5% hydroxylamine to the reaction mixture, vortexing, and incubating at room temperature for 15 minutes. Labeled peptides were then dried via speed vac and reconstituted in 0.1% formic acid to a concentration of 1 µg/µL.

### hFOB Post-Labeling Multiplexing

For full-plex hFOB samples, an equal volume of each of the twelve TMT-labeled hFOB peptide samples was taken and pooled together into one sample. An equal volume of Jurkat peptides, also distinguishable by TMTpro label, was also included in the full-plex pool. For day-plex hFOB samples, separate peptide pools were created for each collection timepoint during hFOB differentiation (0, 2, 4, and 6 days). An equal volume of peptides was taken from each of the three replicates for each day and pooled together, creating a three-plex sample of peptides from one differentiation timepoint. Jurkat peptides were not included in the day-plex samples.

### hFOB Peptide Unfractionated Desalting

Peptide samples intended for unfractionated MS analysis were desalted using Pierce peptide desalting spin columns according to manufacturer protocol (Thermo Scientific 89852). A pH check was conducted for peptide samples prior to desalting to ensure a pH at 3 or below. Solutions with 0.1% trifluoroacetic acid were substituted with 0.1% formic acid. Two 5% methanol washes were conducted immediately before collecting peptide elutions to clear leftover TMTpro reagent. After desalting, samples were dried down and reconstituted in 0.1% formic acid to an estimated 1 µg/µL concentration. Labeled and desalted synthetic peptides were mixed into the desalted endogenous peptide samples (hFOB/Jurkat) at a 2-to-5 ratio of synthetic-to-endogenous peptide by mass.

### Patient-Derived Cell Culture Preparation

Cell populations containing a mixture of mesenchymal stem cells (MSCs), early osteoprogenitors, and committed osteoblasts and osteoclasts^50^ (referred to elsewhere as patient samples or patient-derived MSCs) were collected from the surface of bone shavings collected from four volunteer patients receiving hip replacement surgery. Samples were collected under the IRB HSR protocol # HSR 200300 entitled "Biobank for the systems genetics of bone diseases" approved by the University of Virginia IRB for Health Sciences Research. The original patient identification numbers (9, 14, 16, and 18) are used to distinguish patients in MS raw file names but have been simplified (to patient A, B, C, and D respectively) in figures and written text.

Based on prior genotyping, the expected baseline of mineralization by active osteoblasts was highest for patient 14 and lowest for patient 18. Cells collected from each patient were cultured with three media conditions: basal medium, osteogenic medium, and osteogenic medium supplemented with 100 ng/mL bone morphogenic protein 2 (BMP-2) (Peprotech 120-02). Osteogenic medium was used to induce differentiation and mineralization of osteoblasts within the cell population; supplemented media was used to accelerate the mineralization process^51–53^. Following ten days of initial media treatment, cells were harvested and stored as pellets with approximately 3×10^5^ cells each.

### Patient-Derived Peptide Digestion, Labeling, and Cleanup

Peptides were harvested from patient-derived MSC/osteoprogenitor cell pellets using the EasyPep sample preparation kit (Thermo Scientific A40006). Pellets were lysed via suspension and pipetting in lysis buffer, and protein concentration was estimated via 660nm colorimetric assay (Thermo Scientific 22660 and 22663). The full lysate volume was reduced and alkylated with kit reagents prior to digestion with 10 µg trypsin/LysC for three hours at 37°C. After halting digestion, peptides were desalted using kit cleanup columns and reagents. Peptides were dried down, reconstituted in 100 µL 50mM TEAB, and labeled with TMTpro following the previously stated labeling protocol for hFOB peptides.

### High-pH Reversed-Phase Fractionation

Offline fractionation was conducted using the Pierce high-pH reversed-phase fractionation kit (Thermo Scientific 84868). For the full hFOB multiplex and the full multiplex of patient-derived peptides, 5 µg of each of the 13 peptide samples (12 hFOB time points and replicates, 1 Jurkat sample) were pooled together for a total fractionation input of 65 µg. For day-separated hFOB multiplexes, 20 µg of each hFOB biological replicate were pooled together by individual day for a total input of 60 µg. For clinical peptide samples, 5 µg of peptides from each media treatment for a given patient sample was pooled with 5 µg of Jurkat peptides. Prior to fractionation, labeled synthetic peptides (non-desalted) were mixed with labeled endogenous peptide samples (non-desalted) at a 2-to-5 ratio of synthetic-to-endogenous peptide by mass. Collected fractions were dried via speed vac and reconstituted with 0.1% formic acid to an estimated concentration of 0.25 µg/µL.

### MS Liquid Chromatography

Peptides were loaded onto an Acclaim PepMap 100 trap column (300 μm × 5 mm × 5 μm C18) and gradient-eluted from an Acclaim PepMap 100 analytical column (75 μm × 25 cm, 3 μm C18) equilibrated with 96% Solvent A (0.1% formic acid in water) and 4% Solvent B (80% acetonitrile in 0.1% formic acid). Peptides were eluted into the mass spectrometer at 300 nL/min up to 90% B over a period of 2 hours.

### HPLC Fractionated hFOB Proteomics

Global hFOB proteomics analysis was first conducted using MS data originally collected for the bone long-read sequencing study. Sample preparation protocols, MS acquisition parameters, and search parameters are identical to those published under “Quantitative mass spectrometry-based proteomics” in Methods of Abood et al.^14^. A multiplexed sample of TMTpro-labeled hFOB peptides collected over 10 days of differentiation was fractionated into 21 fractions via offline high performance liquid chromatography (HPLC). Fractions were analyzed sequentially via a real-time search (RTS) MS acquisition method using the described 120-minute LC gradient on a Thermo Fisher Orbitrap Eclipse Tribrid instrument. A table of hFOB peptide samples matched to their corresponding TMTpro reagents is provided in Supplementary Materials. Proteome Discoverer (version 2.4.0.305) and R (version 4.3.2) were employed for quantitative analysis of proteins over the differentiation time course. Peptide spectral matching and protein identification were conducted using the hybrid protein database generated by the LRP pipeline. Relative protein and peptide abundances were measured using TMTpro reporter ion intensities collected in SPS-MS3 scans. Gene ontology analysis of proteins of interest was conducted with ClusterProfiler^54^. Multiple protein isoform expression was determined for each protein-coding gene by considering all detected peptides for each gene/protein group and identifying cases of two or more peptide sequences that could not originate from the same isoform.

### Data-Dependent Acquisition MS Parameters

Data-dependent acquisition (DDA) was conducted on a Thermo Fisher Orbitrap Eclipse Tribrid instrument. Positive ion mode was used with 2.0 kV at the spray source, RF lens at 30% and data dependent MS/MS acquisition with XCalibur (version 4.3.73.11). Full MS scans were acquired in the Orbitrap from 375 to 1500 *m*/*z* with 120,000 resolution. Data dependent selection of precursor ions was performed in Cycle Time mode, with three seconds in between Master Scans, using an intensity threshold of 2×10^4^ ion counts and applying dynamic exclusion (*n* = 1 scans within 30 s for an exclusion duration of 60 s and ±10 ppm mass tolerance). Monoisotopic peak determination was applied, and charge states 2–6 were included for HCD MS2 scans (quadrupole isolation mode; 1.6 *m*/*z* isolation window, normalized collision energy at 30%). The resulting fragments were detected in the Orbitrap at 15,000 resolution with standard AGC target and dynamic maximum injection time mode.

### DDA Data Analysis

Proteome Discoverer (PD, version 2.4.0.305) was used to search raw DDA data for peptide spectral matches. Processing nodes employed by PD included Spectrum Files RC, Spectrum Selector, Sequest HT, Percolator, and Reporter Ions Quantifier. Sequest HT was set to full trypsin digestion with up to two missed cleavages allowed and peptide length between 6 and 144 residues. Precursor and fragment mass tolerance were set to 10 ppm and 0.1 Da, respectively. Allowed dynamic modifications included oxidation (M, +15.995 Da), phosphorylation (S/T/Y, +79.966 Da), N-terminal acetylation (+42.011 Da), methionine loss (-131.040 Da) and methionine loss with N-terminal acetylation (-89.030 Da). Static modifications included TMTpro (N-termini and K, +304.207 Da) and carbamidomethylation (C, +57.021 Da). Concatenated target/decoy strategy was validated with q-value with a strict target FDR of 0.01. The custom hybrid protein database generated by the LRP pipeline was used as the search database. TMTpro reporter ion signals, measured within MS2 spectral matches, were used to distinguish protein/peptide identifications by sample.

### Tomahto Internal Standard Parallel Reaction Monitoring (IS-PRM) MS Parameters

IS-PRM targeted acquisition with hFOB and clinical peptide samples was conducted on a Thermo Fisher Orbitrap Eclipse Tribrid instrument using the 120-minute LC gradient described previously. MS acquisition parameters were set using the Tomahto user interface. Method length was set to 120 minutes, mass tolerance for trigger and target fragment ions was set to 30 ppm, isolation width was set to 1.0, and target close-out was set to reset at the end of each injection. All other default settings were kept, including the use of collision-induced dissociation (CID) ion activation and Orbitrap resolution of 15,000 for both trigger and target MS2 scans.

### Tomahto IS-PRM Data Analysis

Analysis of MS data collected for hFOB and clinical samples with Tomahto IS-PRM acquisition was conducted via the Tomahto user interface. Target MS2 scans that contained at least half of the fragment ions observed in the corresponding trigger MS2 scan and did not exhibit MS3 signal from unused TMTpro reporter ion channels were selected and exported in CSV format. Target peptide detection counts and quantitative analysis of isoform-specific peptides in full-plex samples was conducted in RStudio.

### Logistic Regression

Logistic regression was conducted in R (version 4.3.2). Model fitting was conducted with glm() using a binomial link function. In the Figure 1C model, all unique peptide sequences detected in 21-fraction DDA were included in the detected class; the undetected class was created by down-sampling from theoretical undetected peptides (Figure 1A) using a fixed random seed to achieve a 1:5 detected-to-undetected ratio. In the Figure 5A models, all BMD target peptides were considered, with detection class determined by any successful detection with DDA or IS-PRM, respectively. RNA abundance was calculated by summing the mean CPM of each peptide-associated isoform at each timepoint, selecting the maximum value across timepoints, and conducting a log_10_(x+1) transformation. Proteotypicity was calculated with pFly^44^. Peptide position within the protein sequence was calculated by dividing the peptide’s starting amino acid position by the total protein isoform length. This value was calculated for each isoform encoding the peptide, and the minimum value was used for logistic regression. Isoform specificity status (isoform-specific, isoform-informative, constitutive, single-isoform gene, or gene-nonspecific) was treated as a categorical variable with constitutive as the reference. Because no BMD target peptides were gene-nonspecific or from a single-isoform gene, these categories were not included in the Figure 5A models. Model quality was evaluated via Area Under the Receiver Operating Characteristic Curve (AUC-ROC) and Variance Inflation Factor (VIF). Odds ratios and 95% confidence intervals were derived from model coefficients with the broom package.

### Tomahto IS-PRM MS Parameters for Synthetic Peptide Quality Control

To mitigate false positive target peptide detections, an analysis of labeled synthetic peptides without labeled hFOB sample peptides was conducted with Tomahto IS-PRM. Separate analyses were conducted for each of the three sets of 96 target peptides. First, synthetic peptides labeled with shTMTpro were injected alone.

Second, labeled synthetic peptides were spiked into a sample of unfractionated, unlabeled hFOB peptides and injected together. Both injections were conducted using the same gradient and MS acquisition parameters used for IS-PRM acquisition of labeled hFOB peptide samples. Results were analyzed using the Data Analysis mode in the Tomahto user interface. The number of attempted target MS2 acquisitions was recorded for each experiment. False-positive target MS2 acquisitions were counted for any target peptide that showed a target MS2 spectrum with at least one half of the fragment ions detected in the corresponding trigger MS2 spectrum.

