## Supplemental Text and Figures (1-6) for "Targeted proteomics informed by long-read RNA-sequencing reveals protein isoform expression associated with bone mineral density"

Micah D. Lehe (1)

Vasilii Pavelko (1)

Erin D. Jeffery (1)

Vedavathi Madhu (2)

Larry D. Mesner (2)

Megan D. Schertzer (1)

Abhijit S. Dighe (2,3)

Charles R. Farber (2,4)

Gloria M. Sheynkman (1,4,5)

1. Department of Molecular Physiology and Biological Physics, University of Virginia, Charlottesville, Virginia, 22903, US
2. Department of Genome Sciences, University of Virginia, Charlottesville, Virginia, 22903, US
3. Department of Orthopedic Surgery, University of Virginia, Charlottesville, Virginia, 22903, US
4. Department of Biochemistry and Molecular Genetics, University of Virginia, Charlottesville, Virginia, 22903, US
5. UVA Comprehensive Cancer Center, University of Virginia, Charlottesville, Virginia, 22903, US

**Supplemental Text**	2

**Supplemental Figure 1**	4

**Supplemental Figure 2**	5

**Supplemental Figure 3**	6

**Supplemental Figure 4**	7

**Supplemental Figure 5**	8

**Supplemental Figure 6**	9

**Supplemental Tables 1-7**	XLSX

**SUPPLEMENTAL TEXT**

**Shotgun MS achieves broad osteoblast proteome coverage**

Our MS analysis of the hFOB proteome via HPLC fractionation revealed dynamics of protein expression that reflect expectations for osteoblast differentiation. Proteins upregulated in early differentiation were enriched for cell-substrate adhesion, extracellular matrix organization, and ossification (**Supplemental Figure 1C**), key processes as osteoblasts form new bone tissue and anchor to existing matrix[^55–57^](https://sciwheel.com/work/citation?ids=3592084,3592033,17652005&pre=&pre=&pre=&suf=&suf=&suf=&sa=0,0,0&dbf=0&dbf=0&dbf=0). Proteins upregulated later in differentiation were enriched for chemotaxis (**Supplemental Figure 1C**), consistent with mature osteoblasts migrating and depositing new mineral during bone remodeling[^58,59^](https://sciwheel.com/work/citation?ids=17900910,9483777&pre=&pre=&suf=&suf=&sa=0,0&dbf=0&dbf=0). To our knowledge, our data represents the most thorough proteomic dataset of differentiating osteoblasts reported to date[^60–62^](https://sciwheel.com/work/citation?ids=17389386,17389385,17389401&pre=&pre=&pre=&suf=&suf=&suf=&sa=0,0,0&dbf=0&dbf=0&dbf=0).

**Osteoblast biomarker quantification supports hFOB similarity to *in vivo* bone mineralization**

We evaluated RNA and protein expression of osteoblast differentiation markers to support the relevance of the hFOB model to *in vivo* bone mineralization (**Supplemental Figure 1D**). First focusing on RUNX2, a key transcription factor upregulated early in osteoblast mineralization[^63,64^](https://sciwheel.com/work/citation?ids=1331654,330426&pre=&pre=&suf=&suf=&sa=0,0&dbf=0&dbf=0), we observed RNA expression at its highest level at day 0 of differentiation while protein expression remained low but consistent. We repeated this analysis for two key RUNX2-regulated osteoblast biomarkers, COL1A1 and COL1A2[^63,65^](https://sciwheel.com/work/citation?ids=1331654,63772&pre=&pre=&suf=&suf=&sa=0,0&dbf=0&dbf=0). Though both were expected to increase in abundance during early mineralization, this pattern was only observed at the RNA level. We attributed this to the difficulty of sampling ECM-localized proteins with our peptide preparation method[^66^](https://sciwheel.com/work/citation?ids=14513197&pre=&suf=&sa=0&dbf=0). ALPL, another ECM-localized RUNX2-regulated biomarker, also showed upregulation at both the RNA and protein levels following the start of mineralization. Expression of *ALPL* is most critical during early bone calcification, consistent with the observed decrease in expression observed after four days[^67^](https://sciwheel.com/work/citation?ids=17389348&pre=&suf=&sa=0&dbf=0). Lastly, we observed upregulation of another RUNX2-regulated biomarker, FN1, following peak expression of RUNX2[^63,68^](https://sciwheel.com/work/citation?ids=1331654,3592128&pre=&pre=&suf=&suf=&sa=0,0&dbf=0&dbf=0). We explored RNA-protein correlation for other osteoblast biomarkers, including BGLAP/OC, SPP1, BSP, and DMP1[^63,68^](https://sciwheel.com/work/citation?ids=1331654,3592128&pre=&pre=&suf=&suf=&sa=0,0&dbf=0&dbf=0). However, we did not detect any protein expression for these genes, possibly explained by their localization to the ECM[^63^](https://sciwheel.com/work/citation?ids=1331654&pre=&suf=&sa=0&dbf=0) resulting in low enough concentration to escape MS detection.

**Deep-fractionation MS reveals expression of a novel sQTL-associated isoform**

One sQTL-containing gene with multiple protein isoforms resolved from the deep-coverage dataset was *MICAL1*, a monooxygenase and cytoskeletal regulator[^69^](https://sciwheel.com/work/citation?ids=2804697&pre=&suf=&sa=0&dbf=0). Out of 14 detected peptides, 13 originate from the isoform with the highest RNA expression, while one unique peptide was specific to an alternative isoform generated by an alternative acceptor splice event at the BMD-associated sQTL (**Supplemental Figure 5**). This isoform was not previously characterized in GENCODE, and there are currently no prior observations of this peptide in public MS databases[^70,71^](https://sciwheel.com/work/citation?ids=6451196,13870407&pre=&pre=&suf=&suf=&sa=0,0&dbf=0&dbf=0). The functional significance of this splicing event is also unknown, as the affected protein domain appears to be a disordered region[^72^](https://sciwheel.com/work/citation?ids=17304800&pre=&suf=&sa=0&dbf=0). MICAL1 thus illustrates how integrating MS with disease proteogenomics can reveal previously uncharacterized and potentially relevant protein isoforms. However, the novel isoform peptide was detected with only one peptide spectral match (PSM) across all sample fractions, highlighting the reproducibility challenges that can occur in isoform detection.

**False-positive target detections with IS-PRM are negligible**

We conducted IS-PRM experiments without TMT-labeled hFOB peptides to understand and mitigate the risk of false-positive target peptide detections (**Supplemental Figure 6**). Prior to any targeted analysis of hFOB protein isoform expression, we used IS-PRM to conduct targeted acquisition of BMD-associated peptide targets in samples of shTMTpro-labeled synthetic peptides alone, as well as samples of labeled synthetic peptides combined with unlabeled hFOB peptides. Under these experimental conditions, there should theoretically be no acquisition of target peptide MS2 spectra as the injected samples will not contain precursor ions with m/z values matching expected targets, as no peptides are labeled with TMTpro reagent.

Nearly all synthetic peptides were detected with the shTMTpro mass shift when labeled synthetic trigger peptides were injected alone (279 out of 288 synthetic peptides) or in an unlabeled hFOB peptide matrix (266 out of 288 synthetic peptides) for IS-PRM acquisition, providing additional confirmation of synthetic peptide stability and labeling efficiency. Target MS2 acquisition was attempted for all detected trigger peptides, but only a few targets had scanning attempts that were reported after passing IS-PRM filtering criteria. IS-PRM ultimately reported 38 target peptide MS2 scans in the trigger-only injection and 10 target scans in the unlabeled hFOB injection. After further scrutiny of the reported target MS2 spectra based on the same criterion used for actual hFOB isoform targeting (i.e., target MS2 must contain at least half of the fragment ions detected in the corresponding trigger MS2), we concluded that 12 target peptides produced false-positive detections in the trigger-only injection. However, the spectra produced from these targets presented other evidence that distinguished them as false detections, including high TMT reporter ion signal from unused reagents or a complete lack of reporter ion signal in MS3 scans. Additionally, there were no reported target MS2 scans in the unlabeled hFOB injection that met target detection criteria. This bolstered our confidence that target peptide acquisitions for our experimental osteoblast samples, particularly for low-abundance isoform-specific peptides, could be trusted if we applied the same validation criteria to those results.

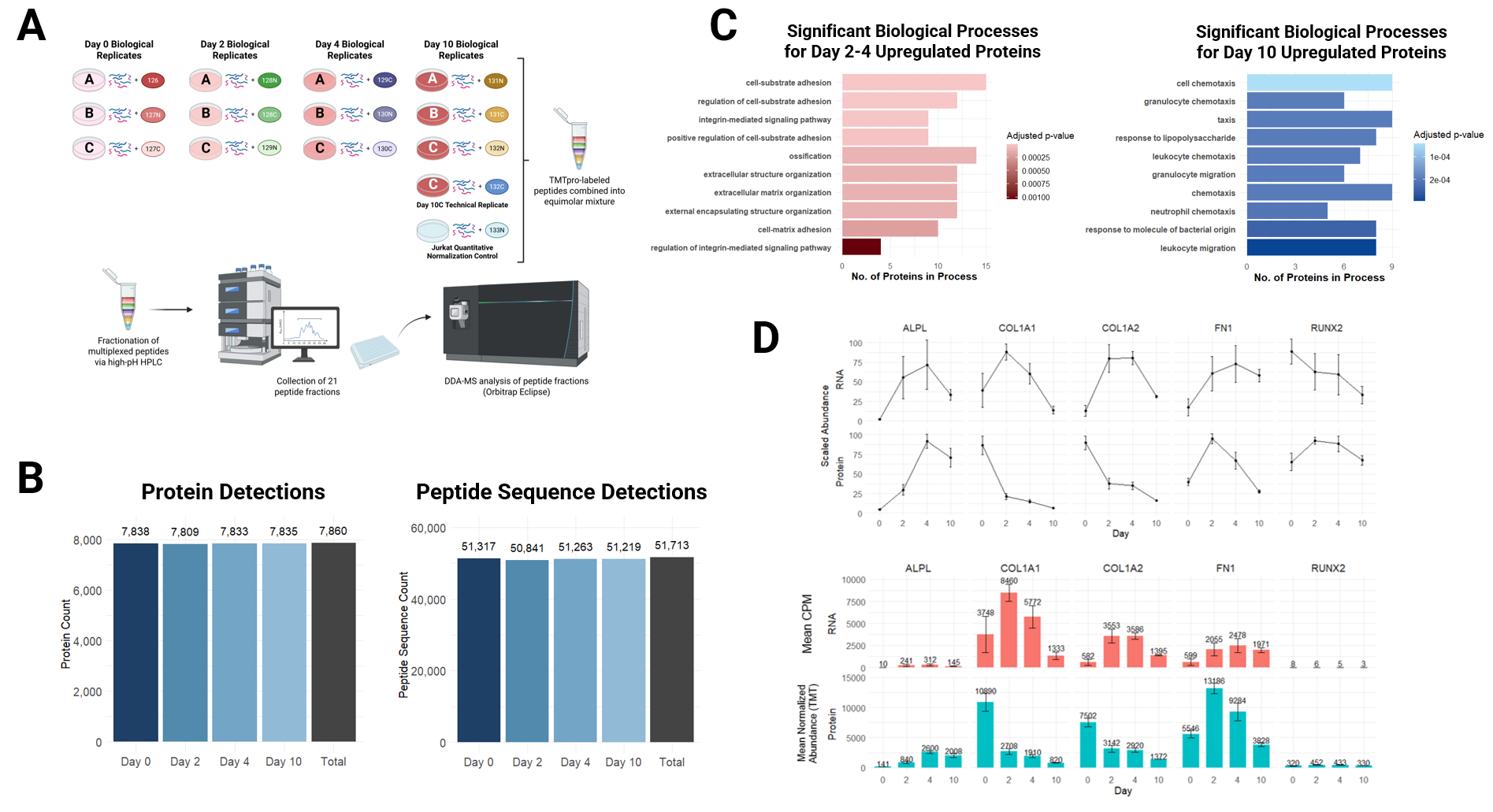

**Supplemental Figure 1: Proteomic profile of deep-fractionation DDA-MS analysis of hFOBs.** A) Schematic of experimental design for MS TMT analysis of differentiating hFOBs (days 0, 2, 4, 10) collected in biological triplicate. B) Number of proteins (not isoform-resolved) and peptide sequences detected in hFOB differentiation timepoints. Peptides with up to two missed cleavages are reported. C) Gene ontology enrichment for protein groups upregulated at Day 2/4 and Day 10. D) (Top) Relative RNA and protein abundance of five selected osteoblast mineralization biomarkers over ten days of hFOB differentiation. Abundances for both RNA (CPM) and protein (normalized TMTpro intensity) are scaled 0 to 100. Error bars denote standard deviation. (Bottom) Mean absolute abundance of selected biomarkers for both RNA (CPM) and protein (normalized TMTpro intensity). Error bars denote standard deviation.

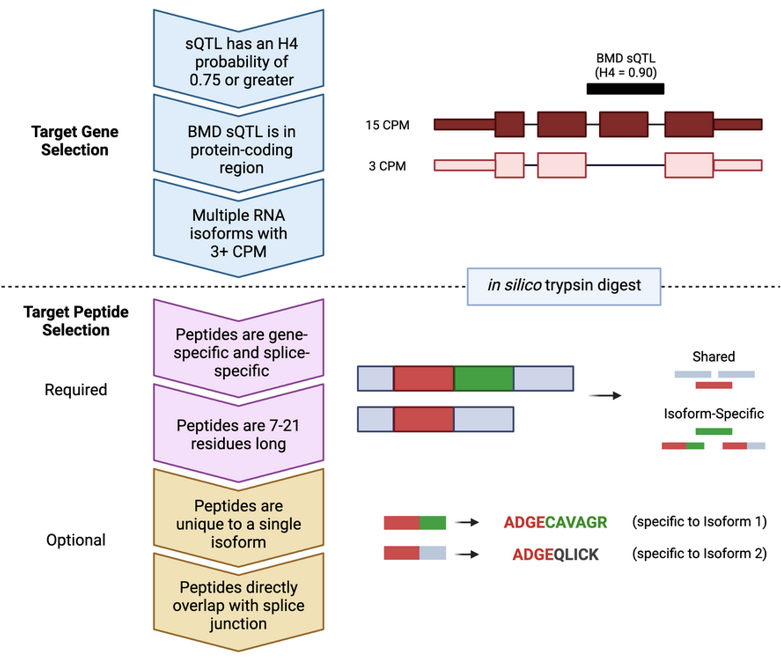

**Supplemental Figure 2: Schematic of target peptide selection process and criteria.**

Top: Target gene/protein groups are selected by prioritizing H4PP score, the presence of a BMD-associated splice junction (sQTL) within the protein-coding region, and the observation of two or more RNA isoforms with a read count of 3 counts per million (CPM) or greater.

Bottom: Protein sequences of selected isoform candidates, determined by open reading frame prediction, are subject to *in silico* trypsin digest with 0 missed cleavages. Candidate target peptides are filtered by length (7-21 amino acids) and gene/protein specificity. Some peptides shared by all or most isoforms within a protein-coding gene group are considered, though priority is given to peptides specific to single isoforms or BMD-associated splice events, especially those that overlap directly with a BMD-relevant splice junction.

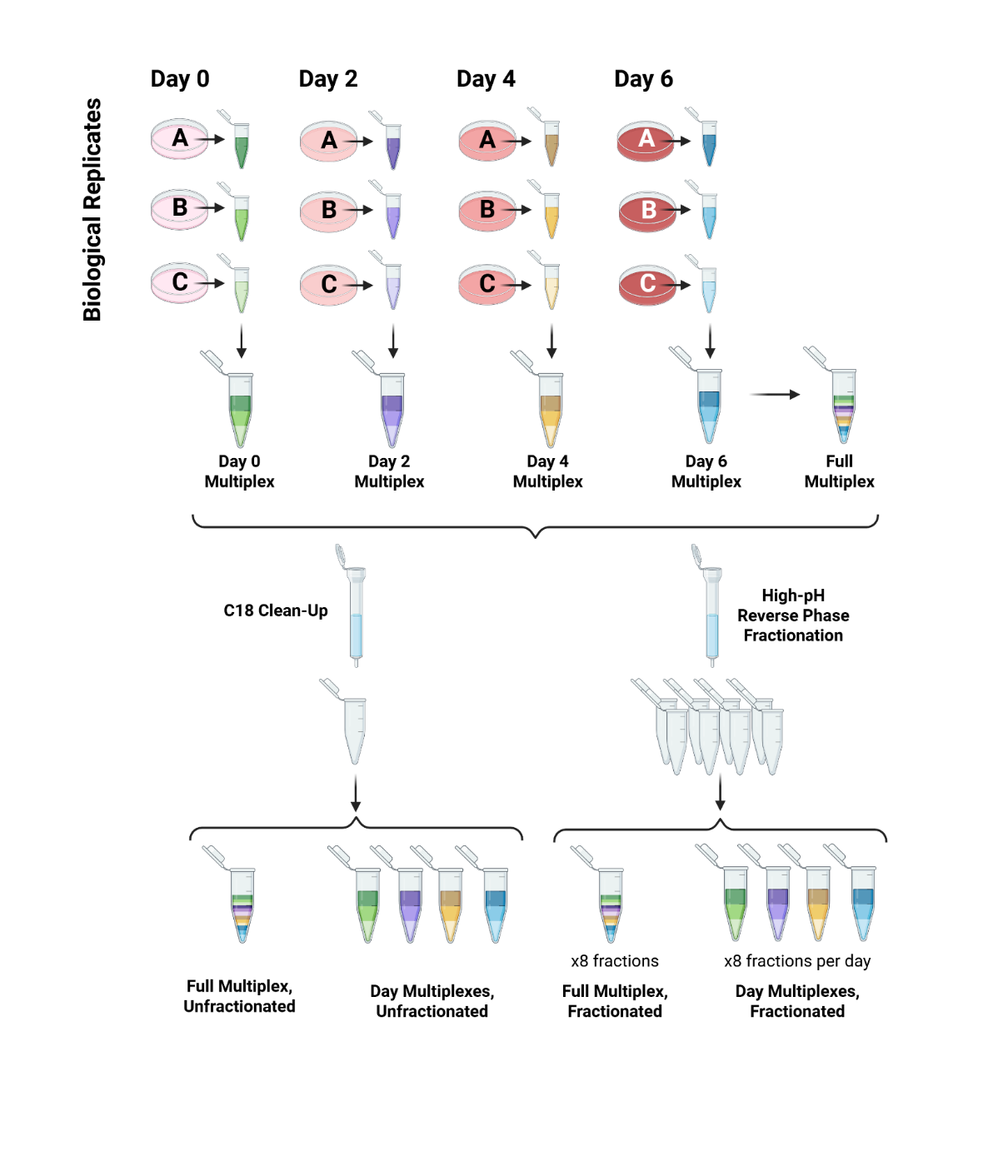

**Supplemental Figure 3: Schematic of peptide multiplexing approach for IS-PRM.**

Top: Human fetal osteoblasts were cultured in 12 replicates, corresponding to four time points of mineralization with three biological replicates each. Peptides collected from each cell sample were labeled with a unique TMTpro reagent and multiplexed together, either separately by day or single pot.

Bottom: Unfractionated peptide multiplexes (left) were desalted via C18 clean-up and analyzed without additional separation. Fractionated multiplexes (right) were desalted via high-pH reverse phase fractionation, generating eight peptide fractions per multiplex.

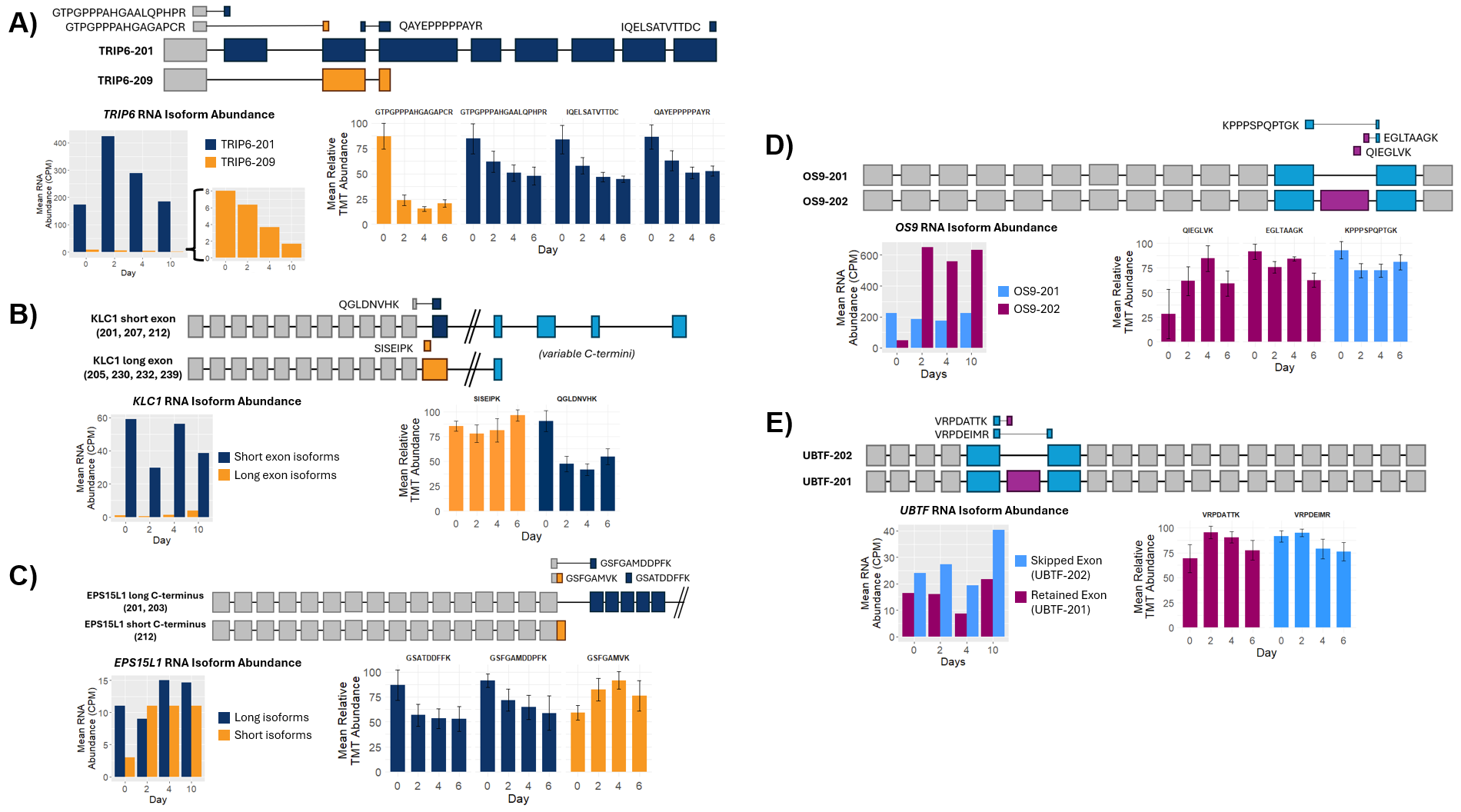

**Supplemental Figure 4: Comparison of RNA and peptide abundance of other BMD target isoforms.** Transcript abundance (RNA counts per million, CPM) of select sQTL-colocalized isoforms and relative TMT-based abundance of target peptides. All protein-coding genes with multiple splice events resolved with IS-PRM are shown, except for *TPM2*. A) RNA abundance and relative peptide abundance for isoforms of *TRIP6.* B) RNA abundance and relative peptide abundance for isoforms of *KLC1.* C) RNA abundance and relative peptide abundance for isoforms of *EPS15L1.* D) RNA abundance and relative peptide abundance for isoforms of *OS9.* E) RNA abundance and relative peptide abundance for isoforms of *UBTF.*

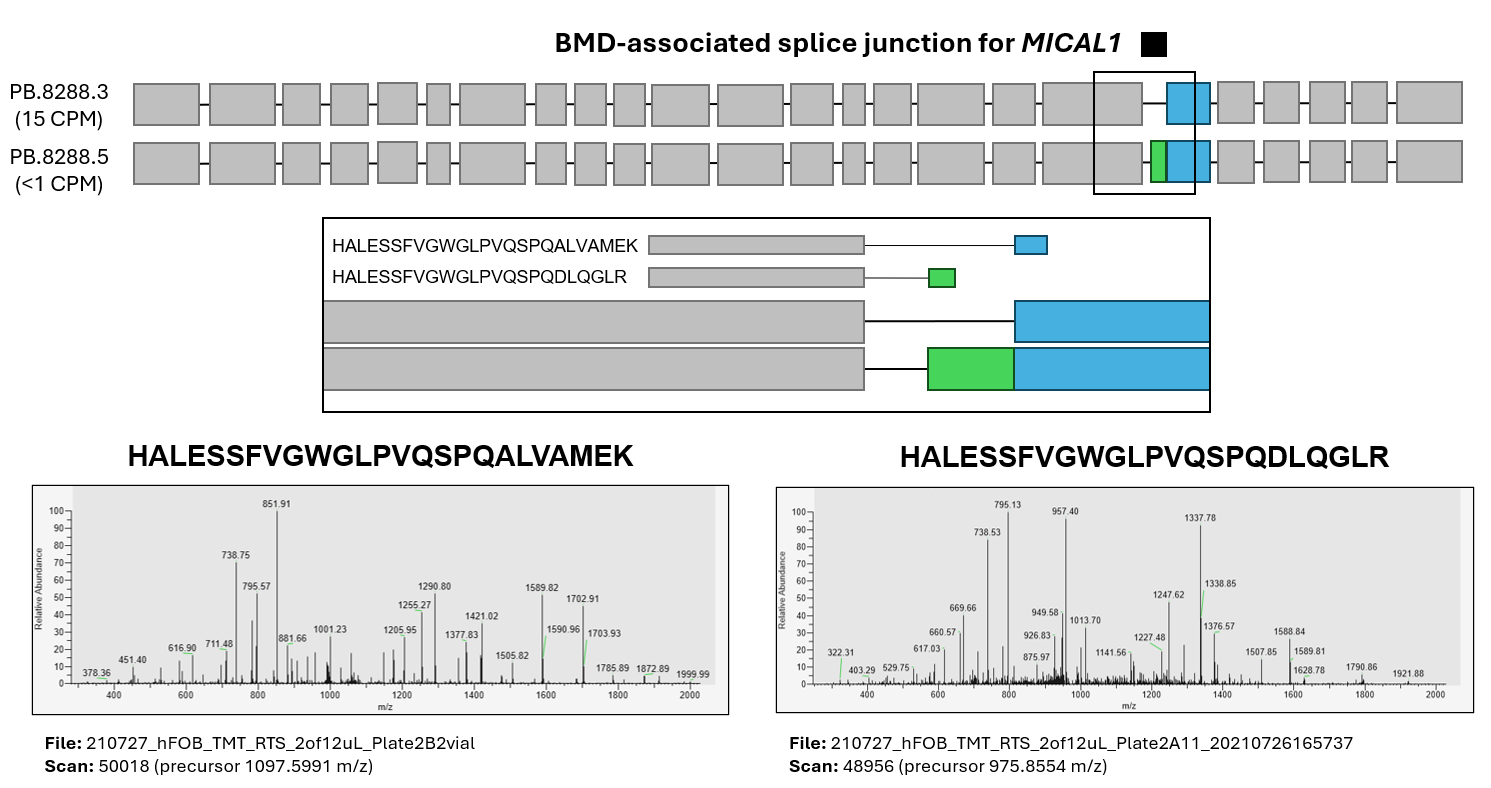

**Supplemental Figure 5: Peptide spectral matches for MICAL1 protein isoforms captured by shotgun hFOB proteomics.** Schematic of MICAL1 isoforms highlights RNA abundance of the major isoform (top) and a minor isoform generated by an alternative acceptor splice event (bottom), overlapping with a predicted BMD-associated splice junction. Mutually exclusive peptides corresponding to the two possible splice events are aligned to the MICAL1 isoforms; peptide spectral matches and scan/precursor information are provided for each.

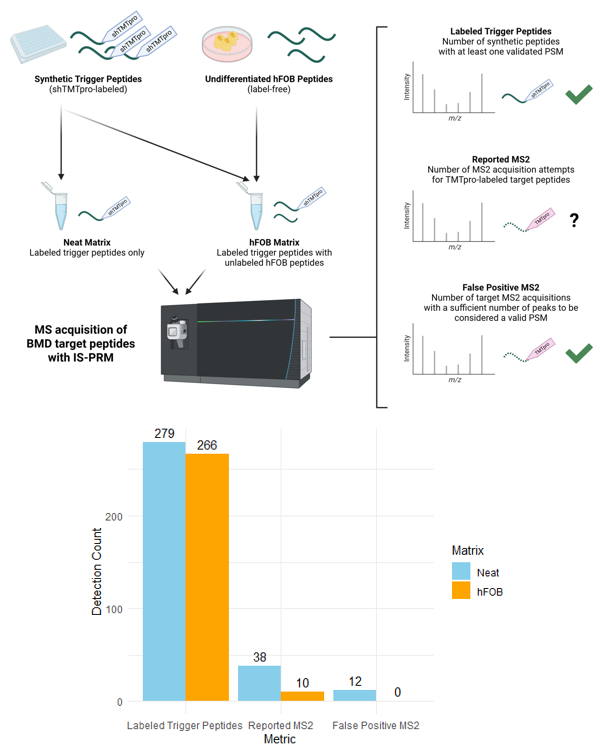

**Supplemental Figure 6: False positive detection of target peptides with IS-PRM.** Targeted acquisition with IS-PRM was conducted with a simple mixture of shTMTpro-labeled synthetic trigger peptides (Neat, blue) and a mixture of shTMTpro-labeled synthetic trigger peptides spiked into a matrix of non-TMTpro-labeled hFOB peptides (hFOB, orange). Total number of labeled synthetic peptides detected, total attempted target peptide MS2 acquisitions, and total number of target MS2 spectra that meet detection criteria are reported on the x-axis.
